# Reliability-weighted target prioritization in CD4+ T-cell Perturb-seq: a generalizability-theory decomposition

**DOI:** 10.64898/2026.07.13.738312

**Authors:** Che Cheng

## Abstract

Genome-scale Perturb-seq screens prioritize candidate targets by the strength of a perturbation’s transcriptional effect. Effect strength does not answer a prior measurement question: is the readout dependable? A large effect estimated from a single guide, a single donor, or a pseudobulk of few cells need not survive replication, and for target prioritization each false lead costs a validation experiment. We treat each perturbation effect as a measurement in a crossed Target × Guide × Donor × Condition design and apply generalizability theory (Brennan, 2001; Cronbach et al., 1972) to separate the dependable part of an effect from facet-specific idiosyncrasy. Guides and donors enter as random facets; condition enters as a fixed facet and is analyzed within its levels. For each target we report a dependability profile over the facets and a joint generalizability coefficient over the two random facets, and we re-rank targets by effect magnitude weighted by that coefficient. On the released screen (Zhu et al., 2025), removing the measurement-error floor estimated from the non-targeting controls raises the number of genes with a dependable target-signal share above .10 from 40 to 7,674. Analyzed within activation states, dependability recovers the T-cell-receptor signaling module as reliably measurable only in activated cells, without recourse to gene annotation. A design study indicates that reliability is limited by the number of guides rather than the number of donors, so a future screen should add guides. Every methodological decision was recorded and adversarially reviewed, and all results regenerate from the released summary statistics.

## 1 Dependability in target prioritization

Genome-scale Perturb-seq couples pooled CRISPR perturbation with a single-cell transcriptomic readout, so that the transcriptional consequence of perturbing each of thousands of genes is measured in one experiment (Dixit et al., 2016; Replogle et al., 2022). Applied to primary human CD4+ T cells, the cells that coordinate adaptive immunity and are a substrate for engineered cell therapies, such a screen identifies the genes whose perturbation most reshapes a T-cell program and, on that basis, nominates candidate targets (Zhu et al., 2025). The nomination carries a cost. Each target that advances is committed to an arrayed knockdown, a functional assay, or an animal model, so a hit that does not reproduce spends a validation experiment on an artifact. What a screen is worth therefore depends not only on the effects it detects but on whether the ranking it induces over targets can be trusted.

Target prioritization conventionally ranks candidates by *effect size*, the strength of a perturbation’s differential-expression signature. The strength is read from the coefficients of a regularized model of expression on guide assignment (Dixit et al., 2016), from a *z*-score or false-discovery rate for each perturbation-gene association, or from the number of downstream differentially expressed genes together with a global test that the transcriptome moved at all (Replogle et al., 2022). Effect metrics have since grown more sophisticated, replacing the gene count with the distance between the perturbed and control single-cell distributions (Peidli et al., 2024) or with a per-target rate of cells classified as perturbed (Papalexi et al., 2021). All of them are measures of effect size, of how large an effect is.

None answers a prior question: whether the effect is dependable, in the sense that it would recur under another guide, another donor, or another cell-state context. The number of differentially expressed genes, for instance, tracks the number of cells assayed (Dixit et al., 2016) and need not track the strength of the perturbation itself (Norman et al., 2019), so a target can rank high on this measure yet low on reproducibility. A large apparent effect can be fragile, when two guides disagree, one donor drives the signal, knockdown is weak, or an off-target flag is present, while a target of moderate but consistent effect is the better validation bet. What decides between them is not which effect is larger in the observed data but which is more likely to reappear when the target is remeasured, because reappearing is what a validation experiment tests. Effect size and dependability can point to different targets, so a ranking that reads only the first keeps nominating the fragile ones. We therefore treat *dependability* as a first-class quantity alongside magnitude, and rank by the two together.

In generalizability theory (Cronbach et al., 1972), the psychometric account of when a measurement can be trusted, dependability has two senses. An effect is dependable when it is *accurate*, a real signal rather than measurement noise, and when it is *generalizable*, recurring across guides, donors, and cell-state contexts. The two show up the same way: an effect that is mostly noise, or that appears only with one guide or one donor, will not reproduce when it is measured again with fresh guides and donors. Measuring that *reproducibility* therefore captures both senses at once. We ask four questions, moving from the present screen to future ones:

- *RQ1 (per target):* which targets carry a dependable effect, one that would recur under a fresh guide and a fresh donor, and does ranking targets by that dependability reorder the shortlist of targets taken forward for validation that effect size alone produces?
- *RQ2 (in context):* within each activation state (Rest, Stim8hr, Stim48hr), does per-target dependability separate a context-specific signal from noise and reconstruct a coherent biological module?
- *RQ3 (the design):* how dependable is the screen as a whole, given its two guides and four donors per target?
- *RQ4 (improving the design):* which facet, guides or donors, limits the screen’s dependability, and how much would a future screen recover by adding to the limiting one?

RQ1 and RQ2 concern individual targets, RQ3 and RQ4 the measurement design, and each answer carries a consequence. A reordering under RQ1 would mean that effect size on its own has been nominating fragile targets and passing over reproducible ones, so the targets a screen sends for validation are the wrong ones: strong but fragile hits crowd out the reproducible ones that would actually confirm. A yes to RQ2 would be strong evidence that the coefficient measures real biology, not an artifact. If dependability reassembles a known signaling module while being told nothing about which genes belong together, the signal it scores cannot be noise, because noise does not organize into a coherent pathway. It would also surface a kind of target that neither effect size nor a pooled ranking can see: one that is dependable in one activation state but not another. RQ3 fixes how far the present screen’s rankings can be trusted, and RQ4 turns that into guidance for the next screen, naming the facet whose reinforcement recovers the most reliability.

At the center of these questions is a per-target coefficient that ranks targets by their dependability. This is not a new concern, and dependability has been pursued along two lines. One measures reproducibility across replicates directly: the irreproducible discovery rate calls discoveries by whether they recur rather than by effect size (Li et al., 2011), and reproducibility metrics tuned to context-specific CRISPR screens show that standard quality-control does not capture whether such a hit will replicate (Billmann et al., 2023). The other decomposes expression variance into named sources, as a per-gene linear mixed model partitions variance into donor, batch, and residual components (Hoffman & Schadt, 2016); data-driven prioritization frameworks likewise integrate several weighted evidence types rather than fitness effect alone (Behan et al., 2019). Neither direction yields a per-target coefficient one can rank on. Generalizability theory joins them: it turns a per-gene variance decomposition into a single dependability coefficient for each target, on the scale of a correlation between independent repeats, and separates the facets a target must generalize across from the context it is measured within.

The rest of the paper develops the per-target coefficient and the two levels it rests on (Section 2), describes the data and the pre-registered choice of estimator (Section 3), and takes up RQ1 through RQ4 in the results (Section 4).

## 2 A generalizability-theory decomposition

### 2.1 The crossed design and its facets

We treat one perturbation’s effect as a measurement and ask how much of it would recur if the experiment were repeated. The quantity of interest is the target gene’s true effect on the transcriptome. What we observe is that effect seen through one guide, in one donor, under one activation state, and read from a finite sample of cells, and each of these can make the observation differ from the truth. This is the setting generalizability theory was built for, and the correspondence is direct (Lord & Novick, 1968): the target gene is the *object of measurement*, its true effect is the *universe score* (the value one would obtain by averaging over infinitely many independent repeats), the *n*_*g*_ guides per target are interchangeable versions of the same perturbation (*parallel forms*), and the *n*_*d*_ donors and the activation states are further experimental factors, or *facets*, crossed with the target, with *n*_*g*_ = 2 and *n*_*d*_ = 4 in this screen. Cell sampling and sequencing contribute the residual noise. Classical test theory writes an observation as a true effect plus error, *X* = *τ* + *ε* (Novick, 1966), and generalizability theory (Cronbach et al., 1972) splits that error into one term per facet,

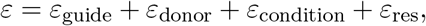

so that the observed variance is a sum of components,

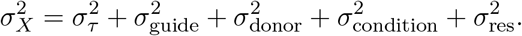

Here 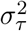 is the variance of the target’s true effect across genes, and 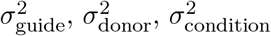 measure how much that effect moves across guides, donors, and states, with 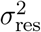 the residual. The analysis estimates each component; these are the variances the coefficients below are built from.

We distinguish random from fixed facets. Guides and donors are *random* facets: one wants the effect to generalize across the guides and donors that were not sampled. Condition is a *fixed* facet: Rest, Stim8hr, and Stim48hr are three deliberately chosen activation states, not a sample from a population of conditions, so the appropriate treatment is a separate analysis within each state rather than a generalization across states. A perturbation that acts only in the activated state is context-specific, not unreliable, and folding condition into a generalization coefficient would misclassify that signal as noise.

### 2.2 Per-target dependability

For target *t*, dependability is a *profile* of three coefficients, one per facet, kept separate rather than multiplied into a single index. Each answers a reproducibility question: with everything else held fixed, how well does the target’s transcriptional signature repeat across a single facet? An observation is a (target, guide, donor, condition) pseudobulk whose effect profile spans the 18,129 genes, measured against a matched non-targeting control. We average a group of observations into a mean profile and read each coefficient off the correlation, across genes, between mean profiles that differ only in the facet of interest. The three share one underlying form, a *signal share*, 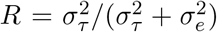, the true-signal variance over the total, where 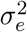 is the error variance of the facet being refreshed; equivalently, its error-to-signal ratio is 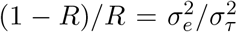. We estimate each one empirically, as a split-half or pairwise profile correlation, rather than as a Gaussian variance-component ratio.^1^

- 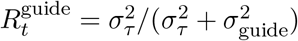: the signal share whose error is the guide variance, estimated by a split-half correlation between the mean profiles of two guide halves; a high value indicates that the signature reproduces across independent guide sets.
- 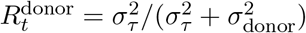: the signal share whose error is the donor variance, estimated by the split-half correlation averaged over the three two-versus-two donor partitions so that it does not depend on donor ordering, then shrunk toward the pooled mean by a moment-based empirical-Bayes step.^2^
- 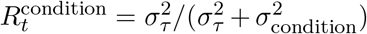: the same signal-share form with the condition variance in the denominator, estimated by the mean of the pairwise correlations among the three within-state profiles. Because condition is a fixed facet, 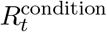 is read as cross-context *consistency* and is kept out of *R*_dep,*t*_: a low value marks a context-specific perturbation, not an unreliable one.

A ranking requires a single number rather than a profile. The per-target dependability *R*_dep,*t*_ provides it, with a concrete interpretation: the correlation expected between two fully independent repeats of the experiment, a second run carried out with fresh guides and fresh donors. Writing an observed profile as a true signal plus independent guide and donor errors, *τ*_*t*_ + *ε*_guide_ + *ε*_donor_, this correlation is again a signal share, now taken over both error sources, 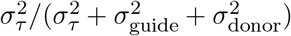. Because the two errors are independent, their error-to-signal ratios add, and *R*_dep,*t*_ follows from the two split-half coefficients:

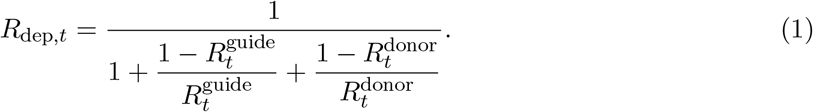

The guide coefficient refreshes one source of variation and the donor coefficient the other; the joint coefficient refreshes both at once, so it poses the most demanding of the three reproducibility questions. The addition of ratios is exact when the guide and donor errors are independent; a guide-by-donor interaction would contribute a further disagreement term, making the assembled *R*_dep,*t*_ an upper bound on the true value.

Dependability measures reproducibility, not importance, so it is not the ranking on its own. We rank by the effect magnitude weighted by dependability,

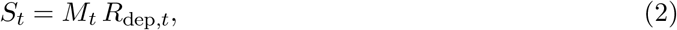

where *M*_*t*_ is the effect magnitude, the root-mean-square of the target’s mean effect profile across genes. A target scores high only when a sizeable effect is also reproducible; a large but fragile effect and a small but reproducible one are both demoted, each for a different reason.

The same construction applies within a single activation state. Restricting the observations to a state *c* ∈ {Rest, Stim8hr, Stim48hr}, we re-form the guide and donor mean profiles from that state’s observations alone, obtain the two within-state split-half coefficients, and combine them through Equation (1) into a within-state dependability 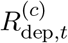. Because each state carries roughly a third of the cells, 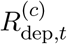 is noisier and runs lower than the pooled *R*_dep,*t*_; a target’s context specificity is therefore read by comparing its 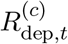 across states rather than against the pooled value.

### 2.3 Design-level dependability

The coefficients so far are per-target. The remaining two questions concern the screen’s measurement design as a whole: how dependable the panel is overall (RQ3), and which facet limits that dependability (RQ4). These are answered at the design level, where the same variance components enter with the design sizes. For a panel of *n*_*g*_ guides and *n*_*d*_ donors, the *generalizability coefficient*, for *relative* decisions such as ranking, is the true-effect share

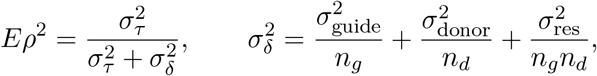

the same guide, donor, and residual variances as above, each now averaged over the design.^3^ The *index of dependability*, 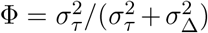, is the coefficient for *absolute* decisions, those that compare a target to a fixed threshold rather than to other targets; its absolute error adds the guide and donor main-effect variances,

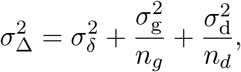

so Φ ≤ *Eρ*^2^. We report *Eρ*^2^ rather than Φ because ranking is a relative decision: those main-effect variances 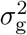 and 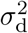, the overall bias of a guide or a donor across targets (as opposed to the within-target variances 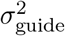 and 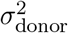, are common shifts that move every target together and leave the ordering unchanged, so counting them as error would understate a dependability that is fully adequate for ranking.

Each coefficient answers one of the four questions. The dependability-weighted score *S*_*t*_ = *M*_*t*_*R*_dep,*t*_ reorders the shortlist relative to effect size (RQ1); the within-state 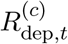, read across states, locates context-specific targets (RQ2); the panel-level *Eρ*^2^ fixes how dependable the screen is overall (RQ3); and the design study, which varies *n*_*g*_ and *n*_*d*_, names the limiting facet and the gain from adding to it (RQ4). The per-target *R*_dep,*t*_ is *Eρ*^2^’s single-target counterpart, the same generalizability read for one target rather than for the panel.

## 3 Data and estimation

The data come from a published genome-scale Perturb-seq screen in roughly 22 million primary human CD4+ T cells (Zhu et al., 2025): CRISPR interference (CRISPRi) silences a single target gene in each cell, and single-cell RNA sequencing reads out that cell’s whole transcriptome. We do not process single cells. We start from the screen’s released *pseudobulk*, the table obtained by averaging all cells that share an experimental combination into one expression profile. Each of its 278,684 rows is one such combination, a silenced target gene, the guide RNA used to silence it, the donor the cells came from, and the T cells’ activation state (Rest, Stim8hr, or Stim48hr). It gives the average expression of 18,129 genes. The screen silences 12,731 target genes with about two guides each, crossed with four donors and those three activation states over two sequencing runs. Only about 45% of that full grid was run, and guides that target no gene serve as the controls. A row averages a median of 49 cells, and as few as one, so rows differ in how precisely they are measured, which is what sets each row’s measurement-error variance. For every row we compute a per-gene effect: the shift in each gene’s expression, in log counts per million, relative to the control cells in the same donor, condition, and run, so every effect is read against controls that share its technical context.

For each gene we ask how much of its perturbation effect is contributed by the guide facet and how much by the donor facet, the two facets a target must generalize across. A crossed mixed model, fit per gene, estimates these variance contributions and weights each pseudobulk by its precision, so rows built from few cells count for less; this is the generalizability-theory design of Section 2. We did not take either estimator for granted. Across the design level and the per-target level we weighed five candidate estimators and chose between them on simulated data whose true variance components are known, in a pre-registered test of which could recover the planted values. At the design level the precision-weighted crossed model was selected. At the per-target level a split-half coefficient with empirical-Bayes shrinkage was the only candidate to recover the planted components under both Gaussian and heavy-tailed noise, and was selected. Of the three set aside, a distance-based decomposition is the clearest case: a share of a pairwise distance is not a variance, and the coefficients are defined from variances. The full estimator decision path (the candidate models and what each can and cannot identify, the two levels of the design, the per-target formulas, and a crosswalk to what the released code computes) is given in Appendix A.

## 4 Results and Discussion

We take up the four questions in turn, moving from individual targets (RQ1, RQ2) to the measurement design (RQ3, RQ4).

### 4.1 Reliability-weighted ranking reorders the shortlist

RQ1 asks whether ranking targets by dependability reorders the shortlist that effect size alone produces. It does. Ranking by *S*_*t*_ correlates with the effect-size ranking at a Spearman *ρ* = .74 over the 7,258 scored targets: the broad order is preserved, but the shortlist itself turns over, with 50 of the 100 highest-effect targets falling out of the 100 most dependable (Figure 1). Reproducible targets that effect size underranks rise in their place (Table 1); RPS3 is the clearest case, 79th by effect size but 4th once weighted by its dependability of .34. These are the targets a dependability-weighted screen would nominate for validation but a strength-ranked screen would overlook. The contribution is a shortlist that effect size alone cannot produce: it demotes strong but fragile hits, which spend validation capacity without confirming, and promotes reproducible effects that a strength ranking buries.

**Table 1:**
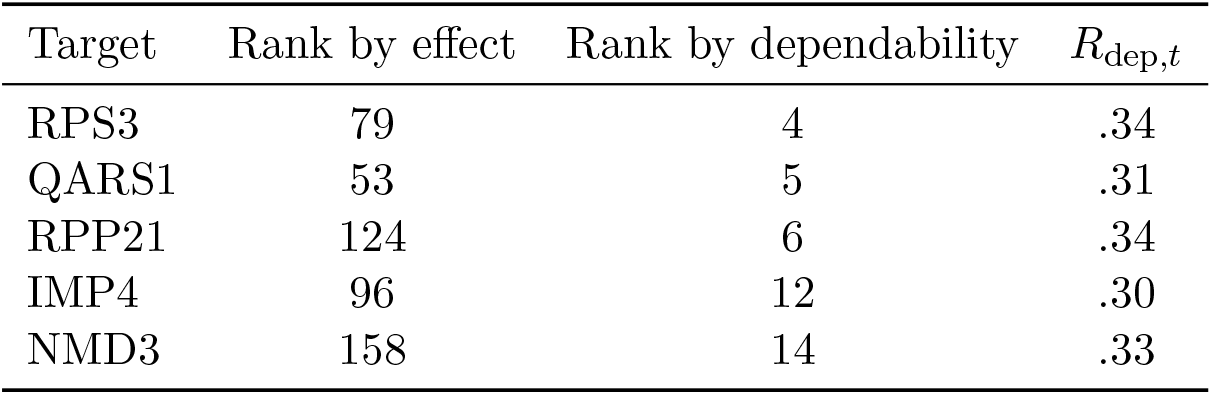
Targets that effect size underranks but dependability promotes. Rank is out of the 7,258 scored targets.

**Figure 1:**
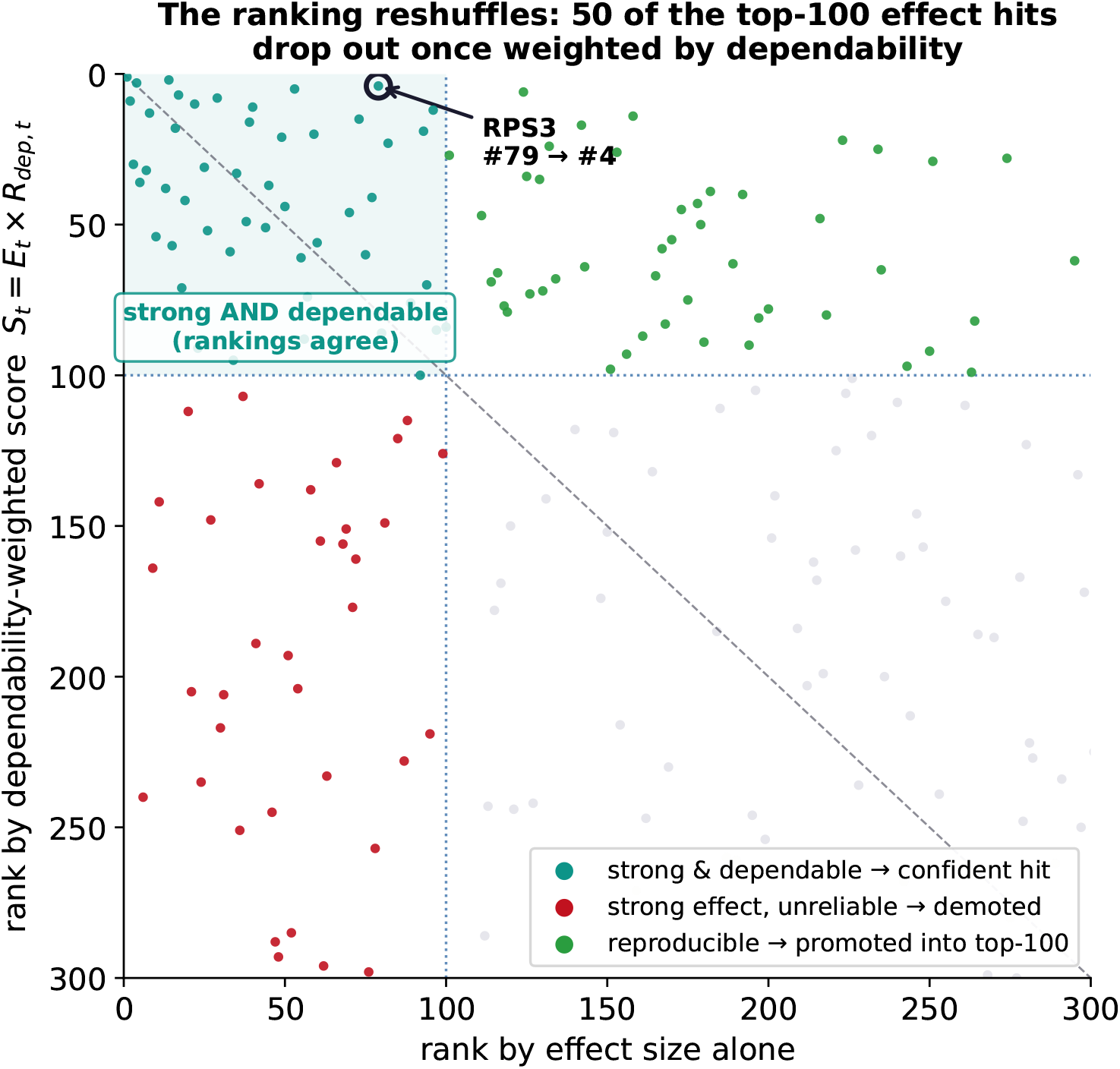
Rank by effect size alone (horizontal) against rank by the dependability-weighted score (vertical); the dotted lines mark the top 100 on each axis. The upper-left quadrant holds targets that are both strong and dependable, where the two rankings agree; lower-left targets are strong but unreliable and are demoted; upper-right targets are reproducible and are promoted. RPS3 is circled.

### 4.2 Context-specific dependability recovers the TCR module

RQ2 asks whether per-target dependability, read within each activation state, separates a context-specific signal from noise and reconstructs a coherent module. Analyzed within activation states, dependability separates targets that are reliable across all states from targets that are reliable in one state and not another (Figure 2). Because a within-state coefficient 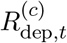 rests on roughly a third of the cells and runs lower than the pooled value, states are compared to one another rather than to the pooled number. Ninety-two targets are context-specific in this sense. Among them, the T-cell-receptor (TCR) signaling module, the CD3 complex (CD3D, CD3G, CD247) with the proximal kinase ZAP70 and the scaffold LAT, is reliably measurable only in activated cells, with resting dependability at the noise floor (resting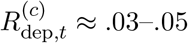; stimulated *≈* .20–.28). The knockdown effect for CD3D grows with activation (effect magnitude 0.26, 0.57, and 1.12 across Rest, Stim8hr, and Stim48hr) and becomes dependable only upon activation. The contribution is direct evidence that the coefficient measures real biology: the framework reconstructs a coherent signaling module from dependability alone, without gene annotation, and noise does not organize into a coherent pathway.

**Figure 2:**
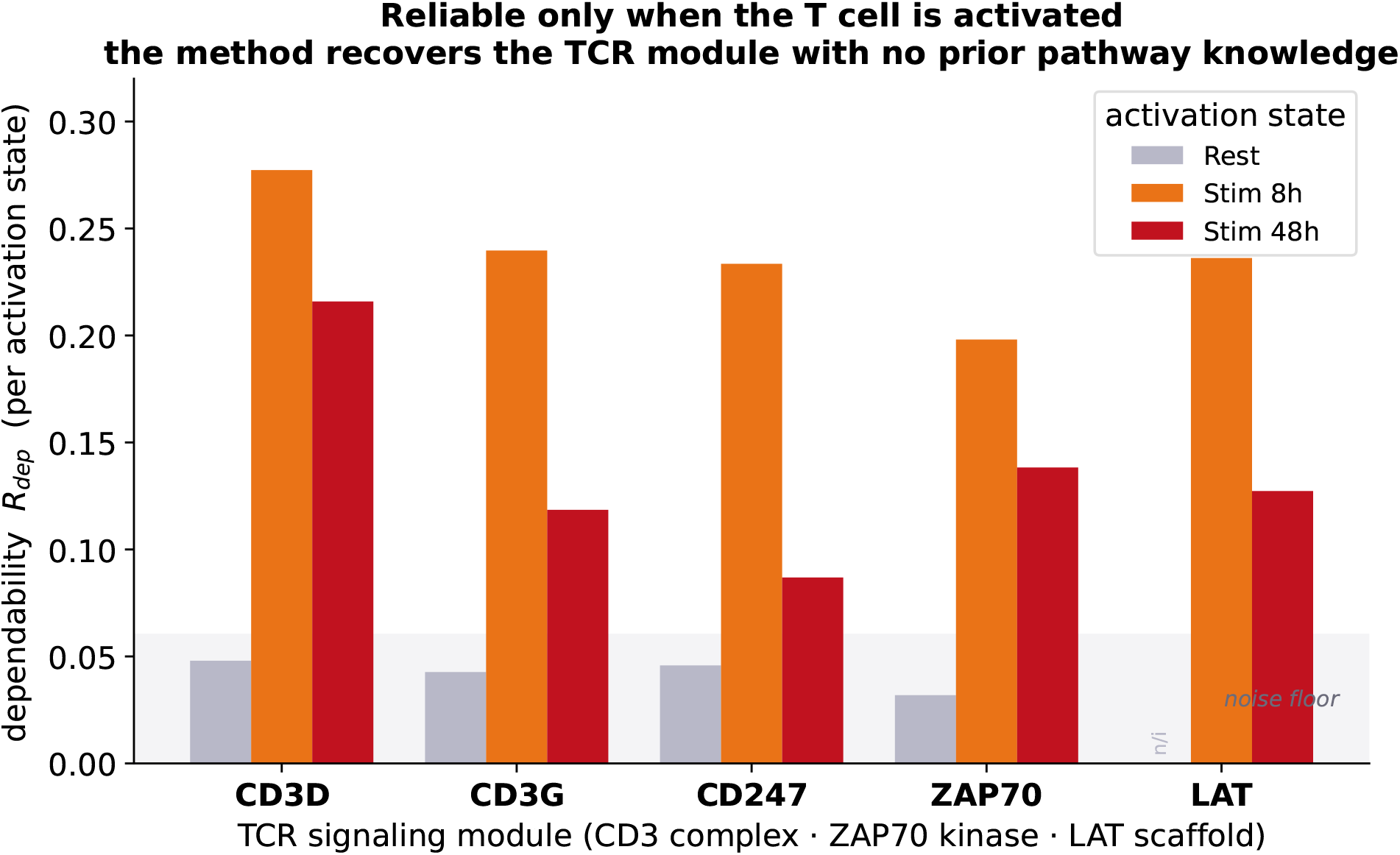
Per-state dependability for the T-cell-receptor signaling module. Resting values sit at the noise floor; the effect becomes reliably measurable only upon activation.

### 4.3 Removing measurement error reveals a moderately dependable screen

RQ3 asks how dependable the screen is as a whole, and answering it means first separating dependable signal from measurement error. The residual of a pseudobulk decomposition combines biological idiosyncrasy with measurement error, and the latter is inflated when a pseudobulk aggregates few cells. The non-targeting controls carry no perturbation, so the variance of the non-targeting effect within its stratum estimates the per-gene measurement-error floor directly. Subtracting this floor from the residual raises the median target-signal share from .019 to .081 and raises the number of genes with a target-signal share above .10 from 40 to 7,674, or 42% of the genome (Figure 3). The dependable signal was present but masked; the same controls give a clean non-targeting-versus-non-targeting negative test (*r ≈* 0), so the correction introduces no signal where none exists.

**Figure 3:**
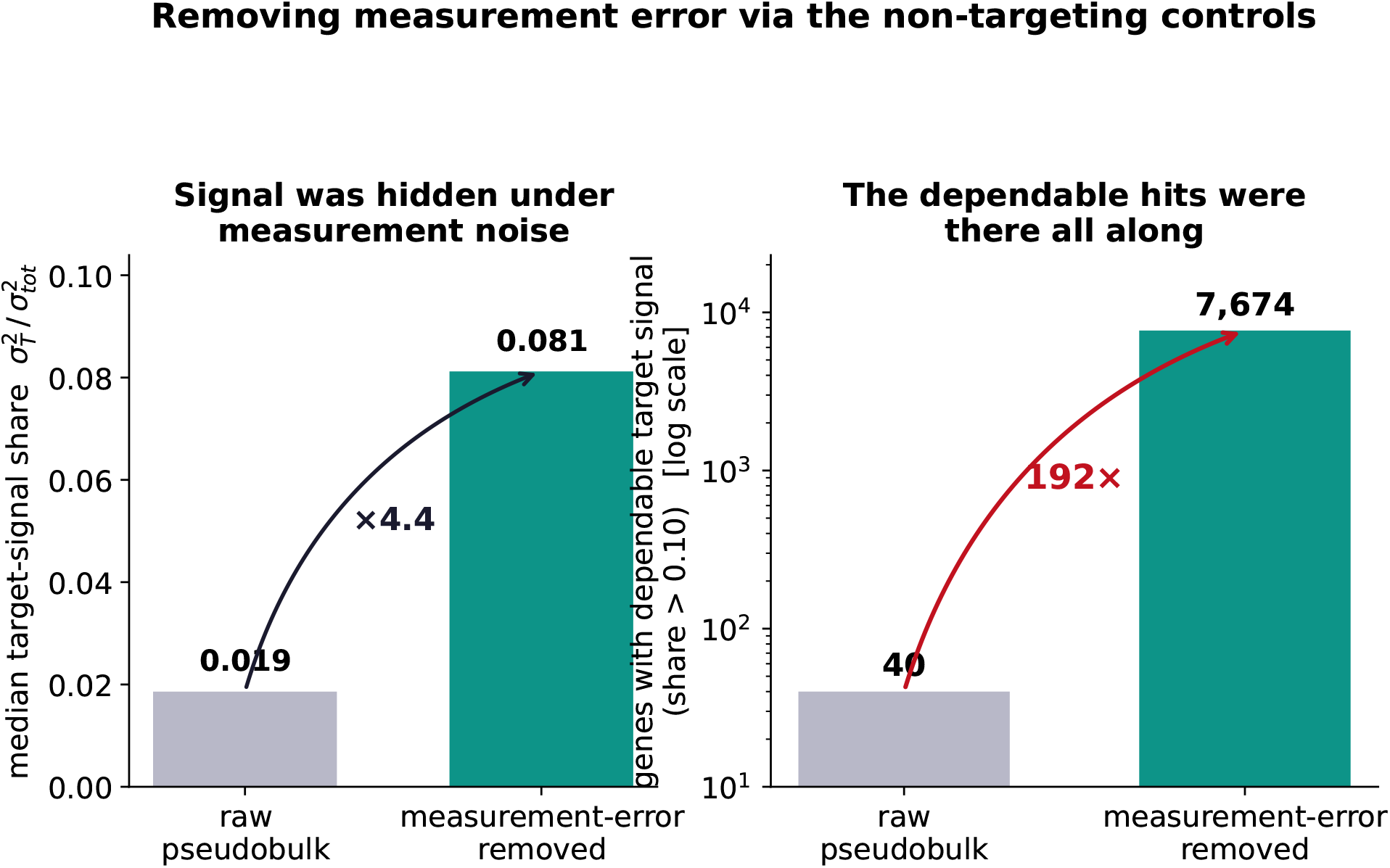
Measurement-error removal via the non-targeting controls. Left, the median target-signal share; right, the number of genes with a dependable target-signal share above .10, on a logarithmic axis.

With measurement error removed, the panel generalizability coefficient for the current design, two guides and four donors, on the representative signal genes is *Eρ*^2^ *≈* .44. Read as a reliability, this is the correlation expected between the target scores this screen produces and those an independent replication with fresh guides and donors would produce, and equivalently the fraction of the between-target variance in the measured effects that reflects real target-to-target difference rather than guide-and-donor sampling error. This is a moderate value: adequate for ranking, where only the order of targets matters, but short of the .70 that an absolute per-target threshold would demand. This calibrates how far the screen can be trusted. Its rankings can be relied on, yet no single target’s dependability clears an absolute bar, so dependability is a prioritization signal for the panel rather than a certificate for any one target read in isolation.

### 4.4 Guides, not donors, limit the design

RQ4 asks which facet limits the screen’s dependability, and how much a future screen would recover by adding to the limiting one. The ranking and its context-specific structure describe the screen as it stands; a design study projects the generalizability coefficient onto alternative numbers of guides and donors (Figure 4). The guide error term exceeds the donor error term by a factor of about 2.4; adding one guide raises the coefficient about three times as much as adding one donor. With two guides the coefficient is capped near .53, and no number of donors reaches .70, whereas .70 is attainable with about 15 guides. Guide-to-guide variability, rather than donor heterogeneity, is the binding constraint, which is actionable guidance for the design of subsequent screens. Repeating the design study within each activation state returns the same verdict in all three states (Table 2), so the recommendation is not an artifact of pooling; within a state the guide term dominates the donor term more sharply still, because most donor disagreement is a cross-state phenomenon. This is actionable guidance for subsequent screens: a fixed budget buys more dependability spent on additional guides than on additional donors.

**Table 2:** Per-condition design study on the representative signal genes. The guide-limited verdict holds within every activation state.

| Activation state | $E\rho^2$ | Guide error | Donor error |
| --- | --- | --- | --- |
| Rest | .502 | 0.026 | 0.005 |
| Stim8hr | .507 | 0.023 | 0.004 |
| Stim48hr | .491 | 0.019 | 0.004 |
| Pooled | .438 | 0.040 | 0.017 |

**Table 3:** What each quantity is in the released code, against its paper-grade target.

| Symbol | Released implementation | Paper-grade target | Status |
| --- | --- | --- | --- |
| $M_{t,c}$ | RMS of the target mean profile | same | exact |
| $R^{\text{guide}}$ | Pearson split-half across genes | (add agreement variant) | consistency only |
| $R^{\text{donor,raw}}$ | two-vs-two split-half, averaged over the 3 partitions | same | exact |
| $w_{t,c}$ | global 0.5 (half-variance split) | Fay–Herriot $\tau^2/(\tau^2 + s_t^2)$ | heuristic; planned |
| $R_{\text{dep}}$ | error-adding joint, fail-closed NA | + interaction term | exact under additivity |
| $S_{t,c}$ | $M_{t,c} \times R_{\text{dep}}$ | same | exact |
| $\sigma_{T,TG,TD,\text{res}}^2$ | REML `lmer` per gene | same | exact (design level) |
| $E\rho^2$ | D-study projection | same | exact |
| me-removal | NTC floor from residual | same | exact (design path) |
| $R_{2g}$ | reported separately | same | exact |

**Figure 4:**
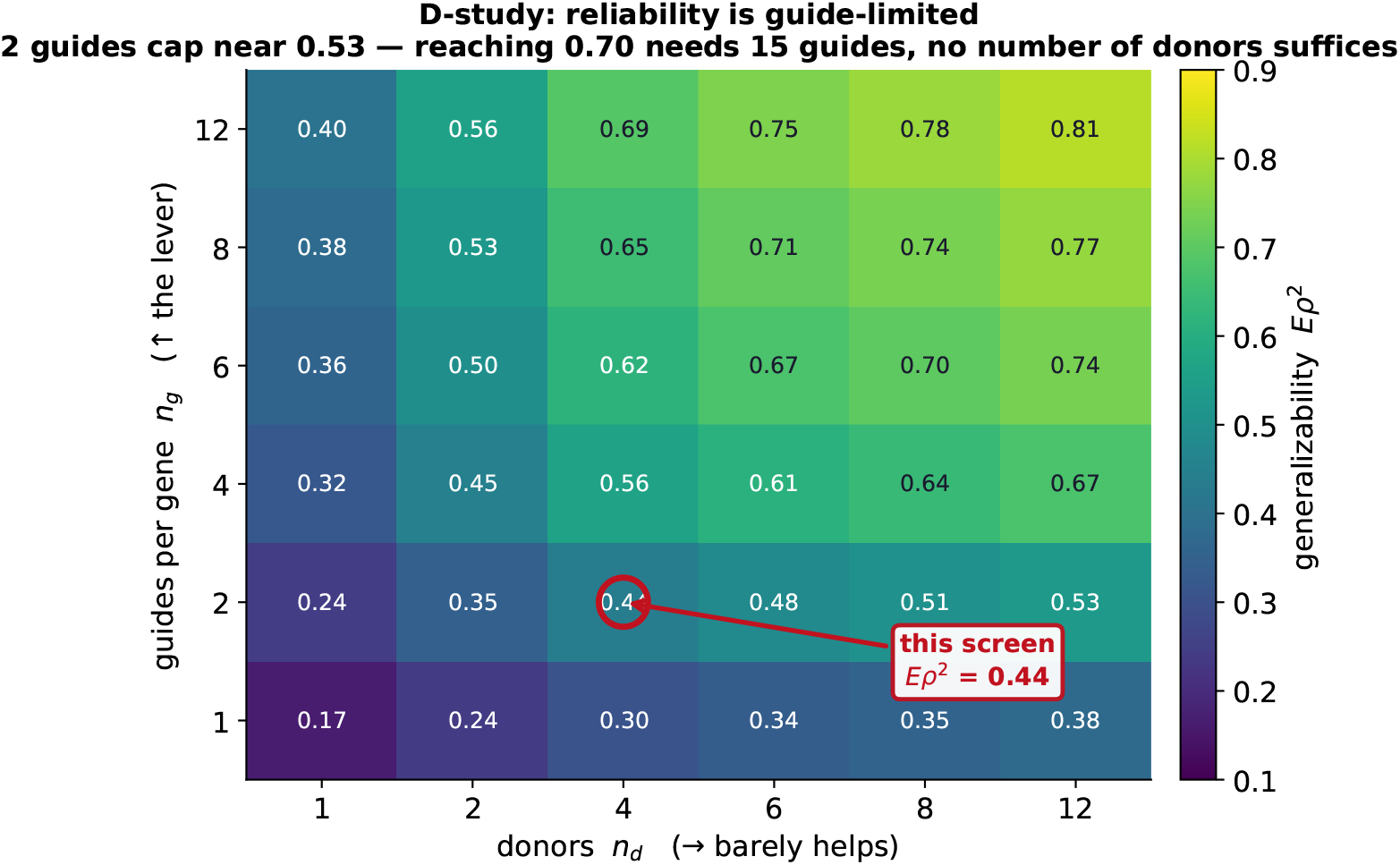
The generalizability coefficient *Eρ*^2^ across the design grid. The current screen (two guides, four donors) is marked; reaching .70 requires about 15 guides and is unreachable by adding donors.

## 5 Conclusions

Generalizability theory gives a genome-scale Perturb-seq screen a per-target dependability coefficient that effect size alone cannot supply. Ranking CD4+ T cell targets by that coefficient reorders the validation shortlist, lifting reproducible effects and demoting strong but fragile ones (RQ1). Within activation states the same coefficient reconstructs the T-cell-receptor signaling module from reproducibility alone, with no gene annotation, which is direct evidence that it tracks real biology rather than an artifact of the summary (RQ2). At the design level the screen is moderately dependable overall, with *Eρ*^2^ *≈* .44 on the representative signal genes (RQ3), and its dependability is limited by guide-to-guide rather than donor-to-donor variability (RQ4), so a future screen recovers the most reliability by adding guides. The analysis needs only the released summary statistics and a pre-registered estimator, regenerates end to end, and records its full decision path for adversarial review. Dependability, not effect size, is the quantity a reproducible screen should rank on.

## 6 Reproducibility

The pipeline runs end to end from the released summary statistics, without processing the full single-cell data: data retrieval, a checksummed evidence manifest, identifiability gates, synthetic method selection, per-gene decomposition, measurement-error removal, the reliability-weighted ranking, the per-context analysis, and the design study. All results are committed as tables and figures, and every reported number regenerates from a single re-run. The code is released under Apache-2.0 at https://github.com/kiki830621/G-perturb.

## 7 Limitations

Four donors leave the donor variance component with three degrees of freedom, so the donor coefficient and the donor axis of the design study carry real uncertainty, which we report as bands rather than point values. Sequencing run is partially confounded with donor, a fact of the released design that we record rather than correct. A small number of full-scale Monte Carlo gate confirmations are scheduled on a compute cluster; the results shown are the local synthetic and real-data tier. The perturbation effect is a plug-in log-CPM contrast; because the pseudobulk is raw counts, we also checked a count-native alternative that scores each effect as a multiplicative fold, a negative-binomial or Poisson log-fold-change with a library-size offset. It recovers a known effect more accurately at low counts on synthetic data, but once low-count genes are handled it leaves the target ranking essentially unchanged, because the plug-in maps a near-zero count to a near-zero effect and so already suppresses the low-count noise that would otherwise inflate sparse targets; the ranking is robust to this choice of effect estimator. Finally, this is a re-analysis of released data, so the design-study guidance is prospective, addressed to subsequent screens rather than to the present one.

## A Estimator decision path

This appendix records the full path from the candidate analysis-of-variance decomposition to the per-target dependability used for ranking, so the estimator can be audited rather than taken on faith. The main text gives the readable version; here we give the design, the identifiability arithmetic, the methods we set aside, the exact formulas, and an honest crosswalk to what the released code actually computes.

### A.1 Estimand, observation unit, and facets

The object of measurement is the target gene *t*. Its response is the effect profile **y**_*tgdc*_ *∈* ℝ^*p*^over *p* = 18,129 genes, the non-targeting-control-relative log-CPM within the matched donor-condition-run stratum. The facets are the two guides per target (nested within target; a random facet we want to generalize across), the four donors (a crossed random facet, with inference confined to the present panel), and the three activation states (a fixed facet whose levels were deliberately chosen, analyzed within level). Sequencing run is partially confounded with donor; the pseudobulk cell count sets the measurement-error variance of each observation, so profiles built from fewer cells are noisier.

### A.2 The crossed model, and why it is not per-target identifiable

Written per outcome gene *j*, the crossed generalizability model is

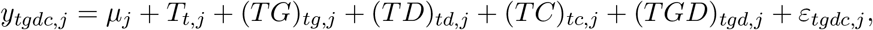

with *T, TG, TD, TGD* random and the condition term *TC* treated as fixed (within-state). The variance components 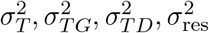 are *design-level* : estimable only by pooling across targets. For a single target in a single condition the design is 2×4 = 8 observations, 7 degrees of freedom: guide contributes 1, donor 3, the guide-by-donor interaction 3, leaving 0 residual. The full 2 × 4 × 3 = 24 factorial is likewise saturated at its highest interaction. Without an identical-specification replicate (a second pseudobulk at the same guide, donor, condition, and run), the top-order interaction and the sampling residual cannot be separated; dropping the interaction to recover a residual is an *additive restriction* the data do not prove, not a demonstration that the interaction is zero. The *≈* 18,000 outcome genes are correlated and do not act as independent replicates. A per-target factorial ANOVA is therefore not identifiable, which forces the two-level treatment below.

### A.3 Two levels, kept distinct

#### Design level

Across targets, a precision-weighted crossed mixed model fit per gene by REML,

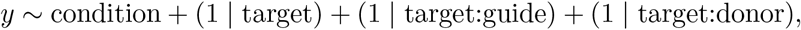

yields 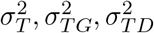 and a residual (the reduced model absorbs the three-way interaction into the residual, per the restriction above). This is an ANOVA / mixed-model variance decomposition. The design study projects it onto alternative designs,

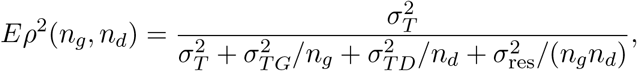

the design-level coefficient of Section 4. Its residual carries measurement error; subtracting the floor estimated from the non-targeting controls is the measurement-error removal of Section 4.3.

#### Per-target level

A single target has only two guides and four donors, so its own variance components are unstable to non-identifiable. We therefore estimate a per-target dependability from split-half profile reproducibility with partial pooling, not from per-target variance components. The two coefficients are not the same quantity and do not share a name: design-level is *Eρ*^2^ (and Φ), per-target is *R*_dep,*t*_.

### A.4 Candidate estimators, and why the others were set aside

The per-target method was chosen by pre-registered synthetic recovery, never by how the real ranking looked. Five candidates were weighed:

1. method-of-moments / factorial ANOVA per target: not identifiable at 2 × 4 (above);
2. precision-weighted crossed REML/GLS: identifiable and used at the *design* level; at the per-target level the same non-identifiability applies;
3. kernel / distance-based decomposition: a distance share is not a variance component without a positive-semidefinite-plus-additivity proof, so it is kept as a diagnostic, not a reported component;
4. robust hierarchical functional model: heavier assumptions and compute, kept as a comparator;
5. per-target split-half correlation with empirical-Bayes shrinkage: distribution-light, identifiable at 2 × 4, and the only candidate to recover the planted components under both Gaussian and heavy-tailed synthetic regimes; selected.

### A.5 The per-target formulas

Within condition *c*, let 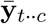 be the target’s mean effect profile. The effect magnitude is the root-mean-square across genes,

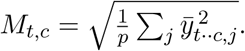

The guide coefficient splits the target’s guides into halves, averages each half’s profile, and correlates them across genes,

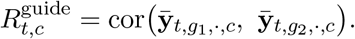

The correlation is Pearson, so it measures *consistency* of the transcriptional pattern, not exact agreement of magnitude; a concordance correlation would also penalize scale and location shifts, and we do not currently apply one. The donor coefficient applies the same split-half construction to donors and averages it over the three balanced two-versus-two donor partitions, so that the raw coefficient 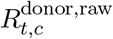 does not depend on donor ordering; it is then shrunk toward the pooled mean *µ*_*c*_,

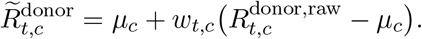

The paper-grade weight is the Fay–Herriot form 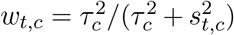 with a target-specific sampling variance 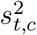. The released code uses a simplified global weight: it sets the within-target variance to half the total variance of the raw donor coefficients, which collapses *w*_*t,c*_ to a single constant *w* = 0.5 applied to every target in the view. The target-varying Fay–Herriot weight is a planned upgrade; the constant is a documented heuristic, not the ideal estimator, and we label it as such.

Writing each coefficient as a signal share 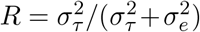, its error-to-signal ratio is 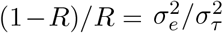; because the guide and donor errors are taken to be independent, the ratios add, and

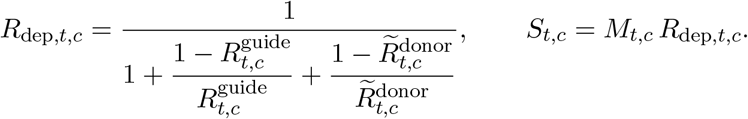

A guide-by-donor interaction would add a further disagreement term, so this form is an upper bound when the interaction is nonzero. If either coefficient is non-positive or undefined the target is reported not identifiable (a fail-closed NA), never zero-filled. The score *S*_*t,c*_ is a ranking criterion, not the probability that a target is real. The deployed screen averages two guides, whose reliability is the Spearman–Brown step-up *R*_2*g*_ = 2*r/*(1 + *r*); because the joint coefficient is defined at the single-guide-by-single-donor level, this step-up is reported separately and is not substituted into *R*_dep_.

### A.6 Pipeline and formula-to-code crosswalk

The analysis proceeds in stages. It begins from the joint pseudobulk, forms each row’s effect and covariance against the matched non-targeting controls, and passes the design through the identifiability and rank gate. The estimator is fixed by the synthetic-recovery comparison; the pipeline then forms the guide and donor split-half profiles, shrinks the donor coefficient toward the pooled mean, and combines the two into *R*_dep_ and the effect-weighted ranking. The same steps repeat within each activation state, and the design study projects the generalizability coefficient onto alternative guide and donor counts. Two paths diverge once the effects are formed. The design-level path carries the variance components, the D-study, and the measurement-error removal; the per-target path carries the split-half coefficients, the shrinkage, and *R*_dep_. Measurement-error removal acts only on the design-level residual, not on the split-half coefficients, which absorb measurement error by construction.

### A.7 Assumptions, costs, and falsifiers

The reported quantities rest on stated assumptions, each of which can fail: guide and donor errors are independent, so a guide-by-donor interaction would make *R*_dep_ an upper bound rather than an equality; the four donors are a panel, not a population, so donor generalization is to these donors; the *≈* 18,000 genes are correlated, so the effective dimension is far below 18,000 and the split-half coefficients are noisier than an independent-gene count would suggest; Pearson measures consistency, not magnitude agreement; the non-targeting-control covariance is shared and heteroskedastic across strata; sequencing run is partially confounded with donor; and without an identical-specification replicate the top-order interaction and the residual are not separable per target. Restoring full per-target variance-component estimation would require more guides, more donors, and lane or library replicates.

## B A recorded, cross-model-hardened process

The methodology was developed under an auditable process (Figure 5). Each decision (the choice of estimator, the thresholds, the classification of each facet as random or fixed, and the criterion for identifiability) was recorded as a versioned issue using issue-driven development (Cheng, 2026a) and spec-driven development, both open-source tooling the author maintains. Because the reasoning was documented rather than implicit, the frozen record could be subjected to adversarial review by a competing frontier model, GPT-5.6 Sol, which returned a blocked verdict with 11 findings, three of them critical, spanning identifiability, measurement error, selection, validation leakage, and compute. Each finding was resolved within the same recorded loop: the falsification gates and controls were frozen before any real result was inspected, the estimator was selected as in Section 3, and a mis-specified reliability aggregation (a product of coefficients) was replaced by the joint coefficient of Section 2.2. The resolutions were then cross-verified across models with a second adversarial pass and with `parallel-ai-agents` (Cheng, 2026b), a package the author maintains that dispatches a task to independent agents and cross-compares their outputs. The recorded decisions, the blocked verdict, and every resolution are versioned in the repository.

**Figure 5:**
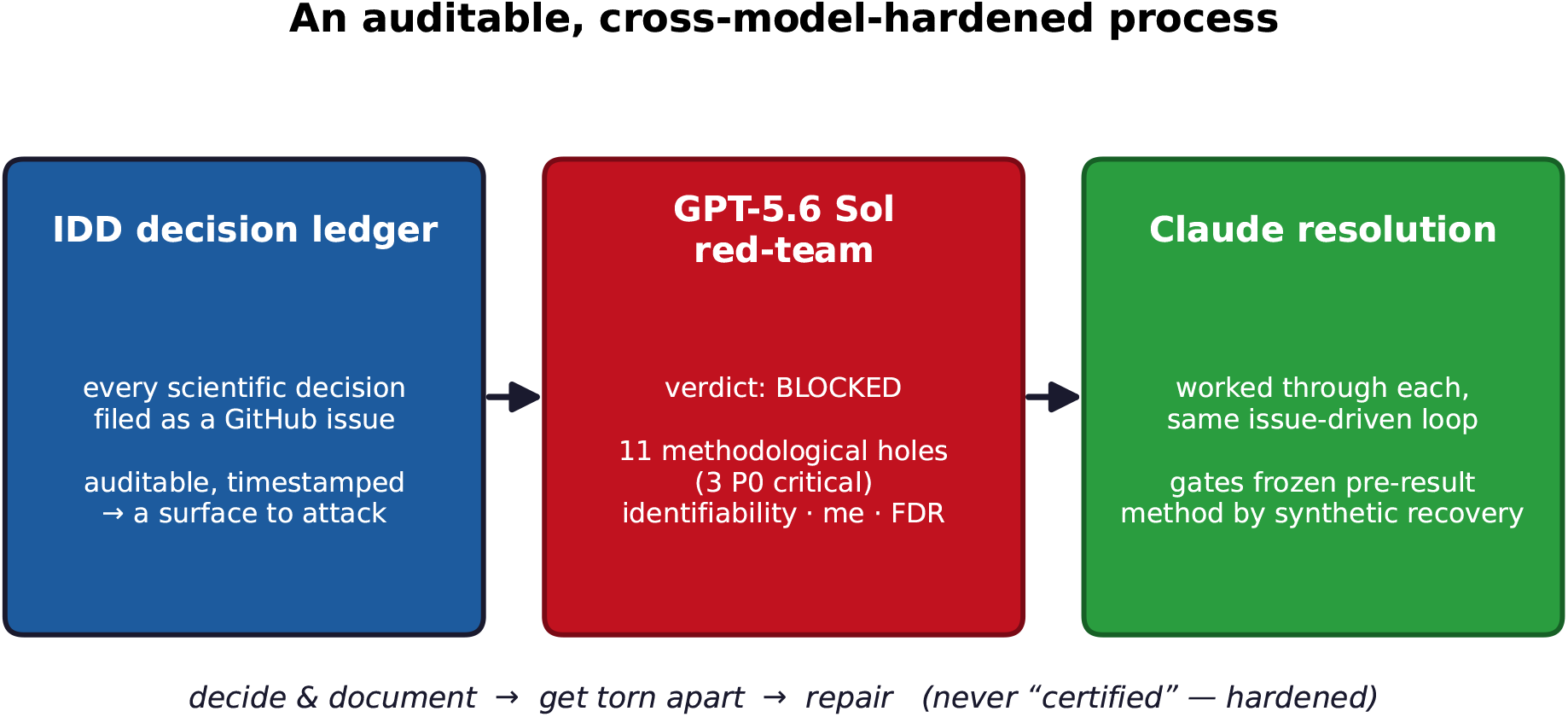
The recorded, cross-model-hardened process: a decision ledger enables adversarial review by a competing model, GPT-5.6 Sol, whose blocked verdict is resolved in the same loop. The verdict was blocked and worked through, not certified.

The manuscript itself was produced under the same discipline. It was drafted and revised in an extended interactive session with Claude Opus, run with a one-million-token context window at maximum reasoning effort, while the author reviewed each section, formula, and claim and directed every revision. The commit history records that revision trail alongside the methodological one, so the writing, like the analysis, is auditable rather than opaque.

## Footnotes

1 These are raw single-observation correlations. For a facet a screen deploys as an average (the two guides, the four donors), the reliability of that average is the larger Spearman–Brown step-up, *kR/*(1 + (*k −* 1)*R*) for *k* units; we report the single-observation form because the joint coefficient below is defined at the single-guide-by-single-donor level. All three are distribution-light empirical correlations, not Gaussian variance-component ratios.

2 With only four donors this coefficient carries three degrees of freedom and is noisy, so it is shrunk toward the pooled mean; the released code applies a global shrinkage factor in place of the target-specific one, as detailed in Appendix A.

3 In the fitted crossed model of Appendix A these are the target, target-by-guide, and target-by-donor components, Written 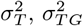, and 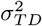.

